# Complete chloroplast genomes of 14 mangroves: phylogenetic and genomic comparative analyses

**DOI:** 10.1101/787556

**Authors:** Chengcheng Shi, Kai Han, Liangwei Li, Inge Seim, Simon Ming-Yuen Lee, Xun Xu, Huanming Yang, Guangyi Fan, Xin Liu

## Abstract

Mangroves are main components of an ecosystem which connect land and ocean and is of significant ecological importance. They are found around the world and taxonomically distributed in 17 families. Until now there has been no evolutionary phylogenetic analyses on mangroves based on complete plastome sequences. In order to infer the relationship between mangroves and terrestrial plants at the molecular level, we generated chloroplast genomes of 14 mangrove species from eight families, spanning six orders: Fabales (*Pongamia pinnata*), Lamiales (*Avicennia marina*), Malpighiales (*Excoecaria agallocha*, *Bruguiera sexangula*, *Kandelia obovata*, *Rhizophora stylosa*, *Ceriops tagal*), Malvales (*Hibiscus tiliaceus*, *Heritiera littoralis*, *Thespesia populnea*), Myrtales (*Laguncularia racemose*, *Sonneratia ovata*, *Pemphis acidula*), and Sapindales (*Xylocarpus moluccensis*). The whole-genome length of these chloroplasts is from 149kb to 168kb. They have a conserved structure, with two Inverted Repeat (IRa and IRb, ~25.8kb), a large single-copy region (LSC, ~89.0kb), a short single-copy (SSC, ~18.9kb) region, as well as ~130 genes (85 protein-coding, 37 tRNA, and 8 rRNA). The number of simple sequence repeats (SSRs) varied between mangrove species. Phylogenetic analysis using complete chloroplast genomes of 71 mangrove and land plants, confirmed the previously reported phylogeny within rosids, including the positioning of obscure families such as Linaceae within Malpighiales. Most mangrove chloroplast genes are conserved and we found six genes subjected to positive or neutral selection. Genomic comparison showed IR regions have lower divergence than other regions. Our study firstly reported several plastid genetic resource for mangroves, and the determined evolutionary locations as well as comparative analyses of these species provid insights into the mangrove genetic and phylogenetic research.

## Introduction

Mangroves are a group of plants that grow on the intertidal zone of the ocean. They are the main and most important component of an ecosystem which connects the land and ocean. Mangroves provide important habitats for marine creatures and benthic organisms and play an important role in regulating energy cycle and maintaining biodiversity [1,2]. These plants have obvious marine adaptations, such as the ability to grow in salty seawater, complex root structures allowing enhanced nutrient absorption and respiratory metabolism, and viviparous reproduction (seeds germinating on trees) [3]. In addition, there are ‘mangrove associates’, also called semi-mangroves, that grow at the edge of the mangroves. Different from the former ‘true mangrove’, these plants are regarded as amphibious plants.

At present, there are approximately 70 species of mangroves, covering 17 families, and 14 mangrove associates from five families [4,5]. Due to their wide distribution, the phylogenetic position of each species would be of considerable interest. As one of the most important organelles in plants, chloroplast has an independent genome with fixed sequence structure and a relatively conserved number of expressed genes associated with energy production and metabolism. In previous reports [6–8], chloroplast genes such as *rbcL* and *psbA* were used to infer evolutionary origins and relationship of mangroves species from different clades or geographical regions. DNA barcodes of *rbcL*, *matK*, *trnH-psbA* genes have also been used to identify unknown mangrove species [5]. To date, however, no study has examined complete chloroplast genome sequences of multiple mangroves.

In this study, we sequenced and assembled the complete chloroplast genomes of 14 mangrove plants (*Pongamia pinnata*, *Avicennia marina*, *Excoecaria agallocha*, *Bruguiera sexangula*, *Kandelia obovata*, *Rhizophora stylosa*, *Ceriops tagal*, *Hibiscus tiliaceus*, *Heritiera littoralis*, *Thespesia populnea*, *Laguncularia racemose*, *Sonneratia ovata*, *Pemphis acidula*, and *Xylocarpus moluccensis*) representing eight families, five rosid orders, and one asterid order. We examined their genome structure, gene content, and simple repeat sequence distribution. In addition, genomic comparative and various molecular evolutionary analyses (phylogeny, selective pressure, genome divergence) were performed to further understand their chloroplast genome characteristics and relationships.

## Results and Discussions

### Chloroplast genome features

A total of 483 Mb sequencing data, with genome coverage ranging from 28× to 526×, were produced for 14 chloroplast genomes in six orders (Table 1): Fabales (*Pongamia pinnata*), Lamiales (*Avicennia marina*), Malpighiales (*Excoecaria agallocha*, *Bruguiera sexangula*, *Kandelia obovata*, *Rhizophora stylosa*, *Ceriops tagal*), Malvales (*Hibiscus tiliaceus*, *Heritiera littoralis*, *Thespesia populnea*), Myrtales (*Laguncularia racemosa*, *Sonneratia ovata*, *Pemphis acidula*), and Sapindales (*Xylocarpus moluccensis*). Using a reference genome-based strategy (see Methods), all chloroplast genomes were assembled into a circular sequence, ranging from 149kb (*Pongamia pinnata*) to 168kb (*Kandelia obovata*) (Table 1, Figure S1). The expected four-segment structure of a plant chloroplast genome was identified in all assemblies: two inverted repeat regions (IRa and IRb), a short single copy (SSC) region, and a long single copy (LSC) region. The length of the four regions were similar within, but slightly different between orders. The average length of IRs of the mangrove orders Fabales, Lamiales, Malpighiales, Malvales, Myrtales, and Sapindales were approximately 23.6kb, 25.6kb, 26.3kb, 26.0kb, 25.3kb, and 27.0kb, respectively. The size of the SSC of the fourteen species ranged from 17.9kb to 20.0kb, while the LSC ranged from 83kb to 91kb (Table 1). The GC content was between 35% to 39%. In IR regions the GC content (~43%) was higher than in SSC (30%) and LSC (34%) regions (Table S1).

**Table 1.**
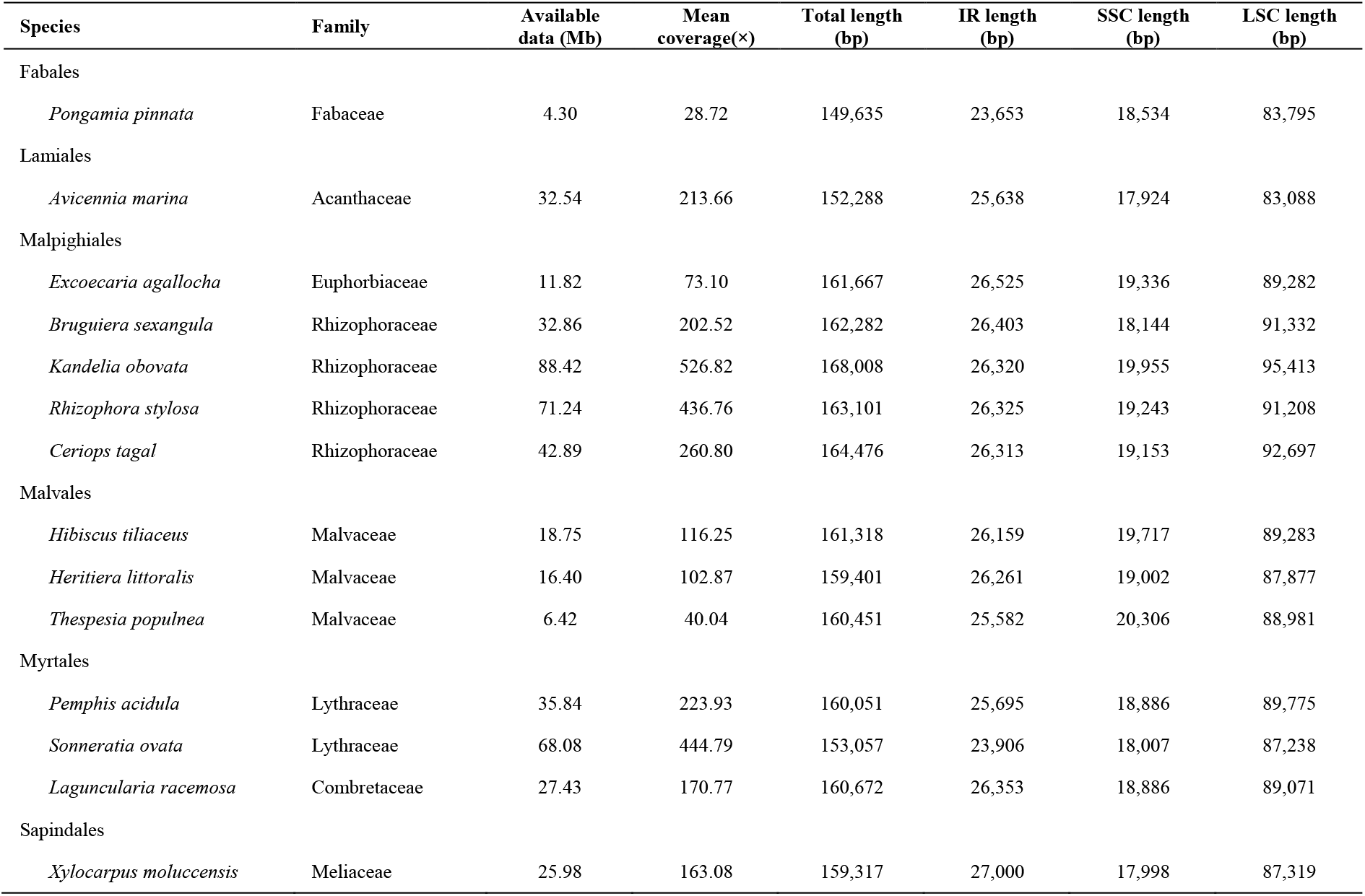
Species information and genome statistics of the 14 mangroves chloroplasts.

The number of plant chloroplast genes is usually highly conserved [9], however there are subtle differences between species [10,11]. The mangrove chloroplast gene number was ~85 protein coding genes, ~37 tRNA genes, and eight rRNA genes (Table 2). The gene component of Photosystems I (five genes), Cytochrome b/f complex (six genes), ATP synthase (six genes), NADH dehydrogenase (12 genes), Rubisco large subunit (*rbcL*), RNA polymerase (four genes), Assembly/stability of photosystem I (*ycf3*, *ycf4*), RNA processing (*matK*), Chloroplast envelope membrane protein (*cemA*), Cytochrome c synthesis (*ccsA*), ATP-dependent protease (*clpP*), fatty acid biosynthetic (*accD*), and Proteasome subunit beta type-1 (*pbf1*) were identical in all the 14 mangrove plastids. Interestingly, *infA*, a chloroplast genome translation initiation factor gene, was only found in *Avicennia marina* and *Heritiera littoralis* (Table 2). This is consistent with a previous study which reported that *infA* is commonly lost in angiosperms, especially in rosids species [12].

**Table 2.**
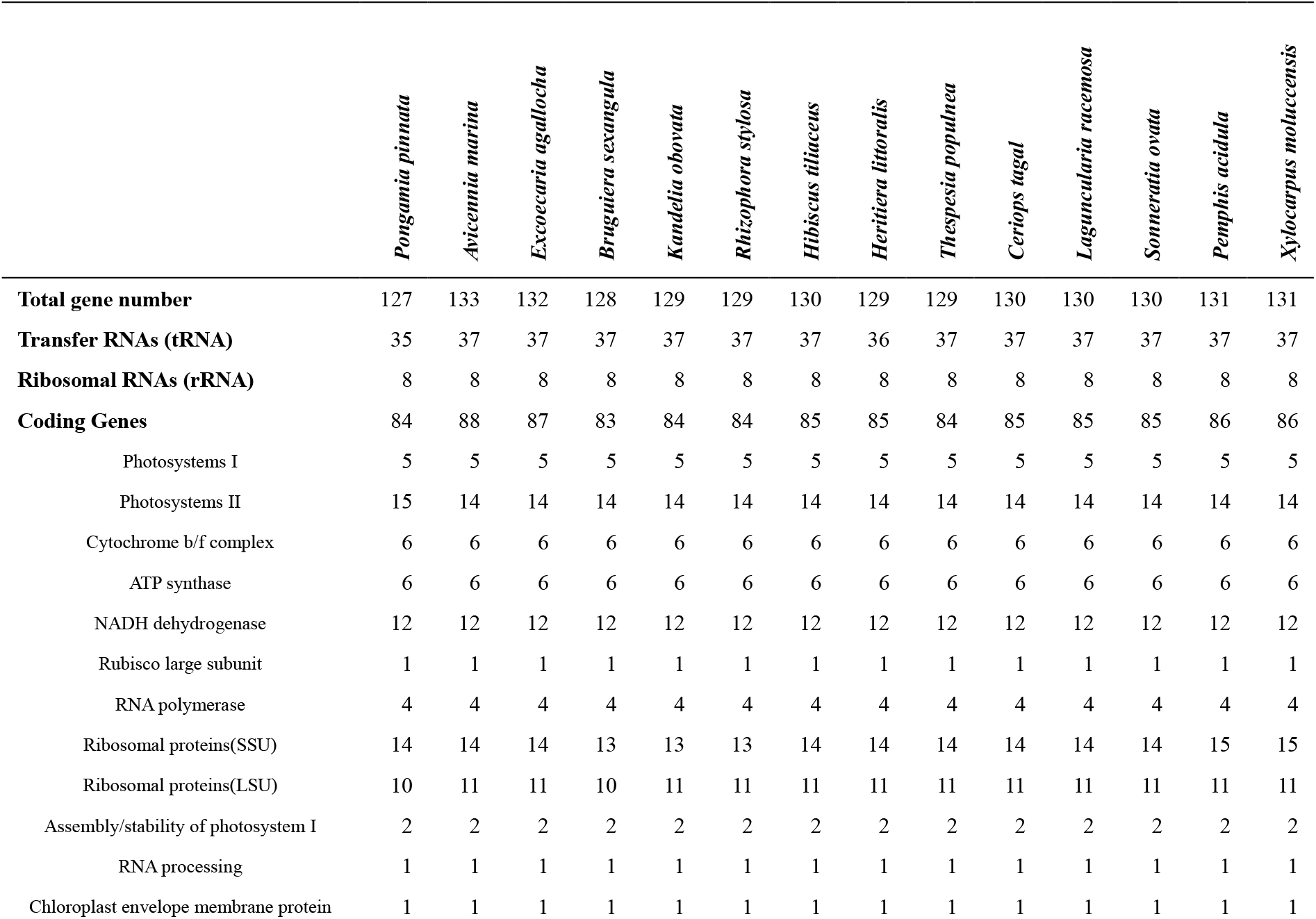
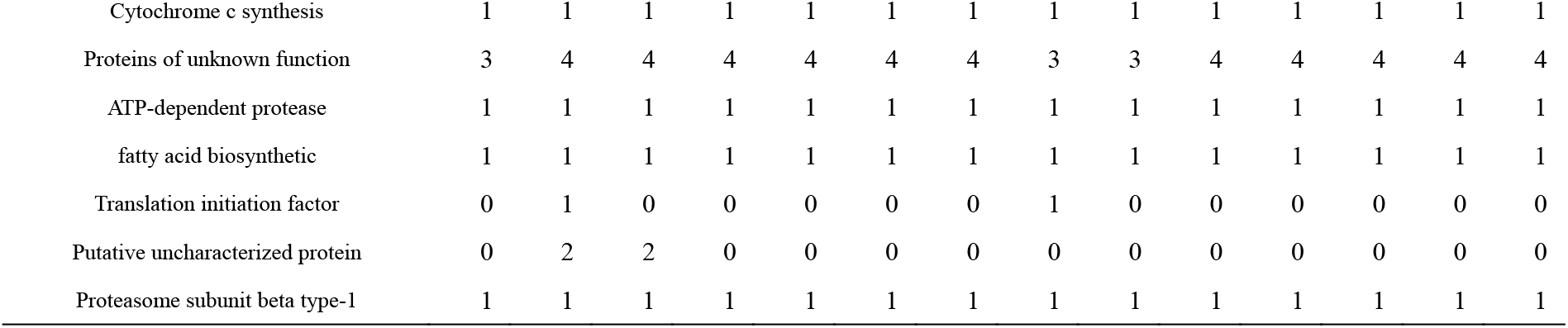
Gene number and function distribution in the 14 mangrove chloroplast genomes.

### Simple Sequence Repeat contents

Simple Sequence Repeat (SSR)s is tandem repetitive (1~6bp) elements and widely distributed in plant plastid genomes. They have been importantly applied as molecular markers for population genetic research and crop breeding because of their abundant genetic diversity [13–16]. In this study, SSRs were detected spanning the 14 mangrove chloroplast genomes. Unlike the chloroplast genes, the SSR content varied widely among species. For example, 130 SSR elements were identified in *Pongamia pinnata*, while only 61 were found in *Avicennia marina* (Table 3). Order Malpighiales (range 133 to 194; *Excoecaria agallocha*, *Bruguiera sexangular*, *Kandelia obovata*, *Ceriops tagal*, and *Rhizophora stylosa*) had a larger SSR number than species in order Malvales (range 80 to 110; *Hibiscus tiliaceus*, *Heritiera littoralis*, and *Thespesia populnea*) and Myrtales (range 88 to 118; *Laguncularia racemosa*, *Sonneratia ovata*, and *Pemphis acidula*) (Table 3). In all mangrove chloroplast genomes, it is found the number of mononucleotide SSRs accounted for at least half of all SSRs (in *Laguncularia racemosa* and *Avicennia marina* up to 80%). Of these, A/T tandems was the most frequent type, followed by dinucleotide, tetranucleotide, trinucleotide, pentanucleotide, and hexanucleotide repeats. These new various SSR data would provide useful resource for future molecular marker analyses.

**Table 3.** Number of SSRs in the 14 mangrove chloroplast genomes.

| Species | Total | u1 | u1:A/T | u1:C/G | u2 | u3 | u4 | u5 | u6 |
| --- | --- | --- | --- | --- | --- | --- | --- | --- | --- |
| *Pongamia pinnata* | 130 | 84 | 82 | 2 | 31 | 6 | 7 | 1 | 1 |
| *Avicennia marina* | 61 | 49 | 45 | 4 | 1 | 4 | 7 | 0 | 0 |
| *Excoecaria agallocha* | 133 | 92 | 87 | 5 | 19 | 7 | 13 | 1 | 1 |
| *Bruguiera sexangula* | 175 | 106 | 103 | 3 | 31 | 14 | 18 | 5 | 1 |
| *Kandelia obovata* | 194 | 106 | 106 | 0 | 36 | 20 | 20 | 10 | 2 |
| *Rhizophora stylosa* | 169 | 115 | 111 | 4 | 22 | 13 | 8 | 9 | 2 |
| *Ceriops tagal* | 142 | 78 | 75 | 3 | 16 | 20 | 23 | 4 | 1 |
| *Hibiscus tiliaceus* | 86 | 55 | 53 | 2 | 15 | 4 | 10 | 2 | 0 |
| *Heritiera littoralis* | 110 | 76 | 74 | 2 | 7 | 6 | 13 | 6 | 2 |
| *Thespesia populnea* | 80 | 47 | 44 | 3 | 17 | 3 | 7 | 5 | 1 |
| *Sonneratia ovata* | 112 | 77 | 72 | 5 | 15 | 5 | 10 | 4 | 1 |
| *Pemphis acidula* | 88 | 64 | 64 | 0 | 7 | 7 | 9 | 1 | 0 |
| *Laguncularia racemosa* | 118 | 98 | 96 | 2 | 6 | 5 | 8 | 1 | 0 |
| *Xylocarpus moluccensis* | 98 | 72 | 71 | 1 | 6 | 6 | 11 | 3 | 0 |

\*u1, u2, u3, u4, u5, and u6 represent unit of mononucleotide, dinucleotide, trinucleotide, tetranucleotide, pentanucleotide and hexanucleotide.

### Phylogenetic relationships of mangroves

Similar to mitochondrial genomes used in vertebrate genetics, chloroplast genomes are highly useful in resolving phylogenetic and evolutionary questions [11]. In order to unfold the phylogenetic relationships of these mangroves, a total of 57 reported chloroplast genome data which represent 57 distinct terrestrial plant families in 17 orders, together with our 14 mangrove species, were used to construct phylogenetic trees (Table S2, Figure 1). Based on the whole chloroplast genome-wide sequences, we identified 41 highly conserved genes (see method) in the 71 species and constructed two Bayesian inference (BI) trees and three Maximum likelihood (ML) trees using different data sets and models (Figure 1, Figure S2-S5). We compared the five evolutionary trees in two ways and found that the trees constructed using whole gene sets have relatively higher posterior probability or bootstrap, and the position of species also tends to be consistent with low log-likelihood difference value between them (Tables S3). It is additionally found that the tree constructed with Bayesian algorithm with 41 gene sequences showed the highest reliability (Table S3) and most branches have posterior probabilities more than 0.9 (Figure 1). These results suggest high confidence of the BI phylogenetic tree in this study.

**Figure 1.**
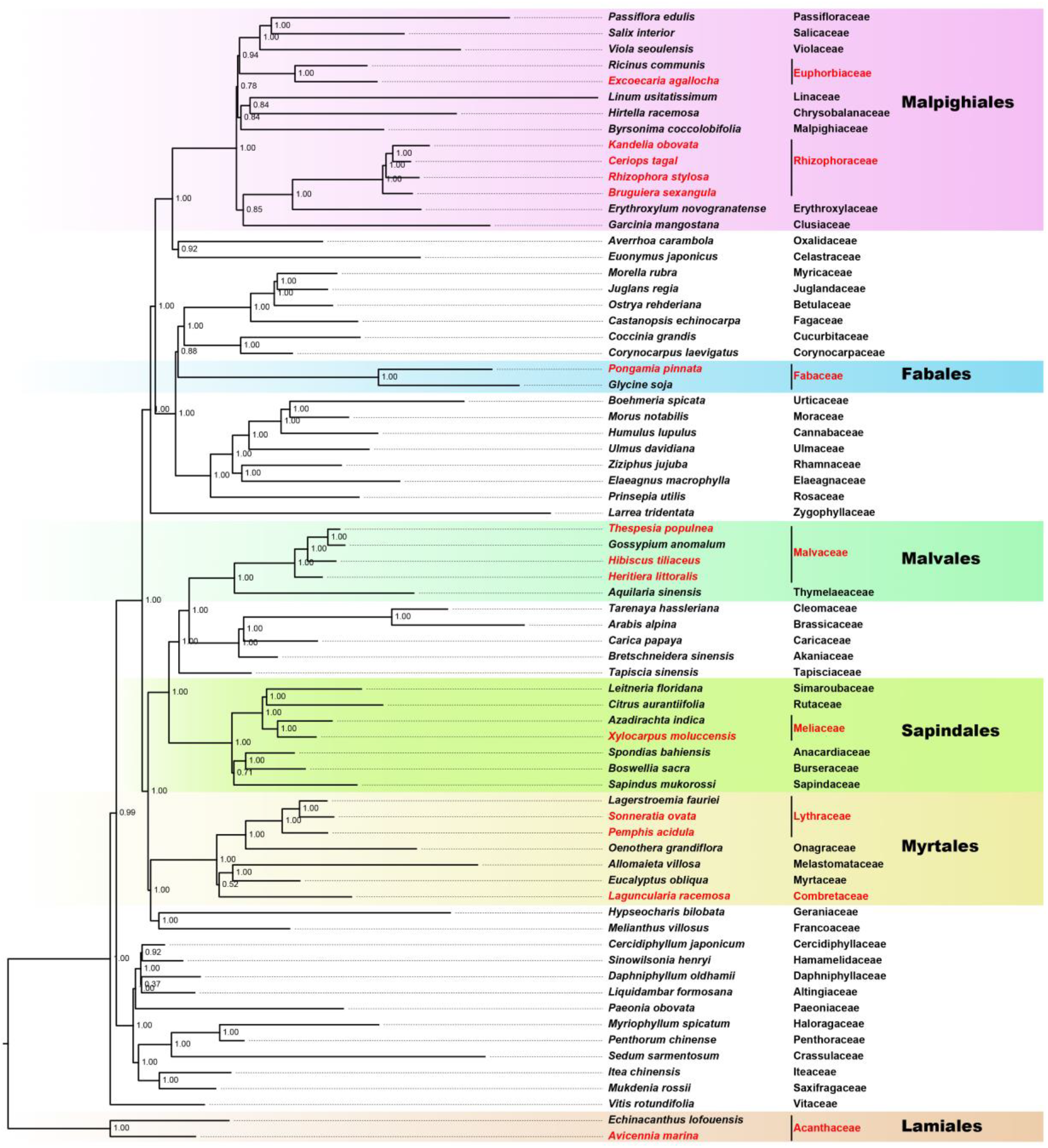
The Bayesian inferred phylogenetic tree based on whole chloroplast genes of 14 mangroves and 57 land species. Mangroves are in red, and the highlight background mark the orders of mangroves and their sister species.

From our phylogenetic tree, at the scale of dicots, it is clear that species in same order are in a group. Mangroves *Avicennia marina*, an asterids, is unsurprisingly close to another asterids plant *Echinacanthus lofouensis*. Using these species as ‘root’, we obtained a tree that distinctly display the phylogenetic relationship of the 13 rosids mangroves. Myrtales is close to Geraniales species. Myrtales here contains five families, in which Myrtaceae and Melastomataceae are in one clade, while Onagraceae and Lythraceae (including two mangroves *Sonneratia ovata* and *Pemphis acidula*) are in another clade, and Combretaceae (*Laguncularia racemose*) is a separate node. The relationship of families within this order is consistent with a ML tree of Myrtales constructed by Berger and colleagues [17]. Sapindales includes one mangrove *Xylocarpus moluccensis*, and their positions in this order here support the result in Toona plastid analysis [18]. For Malvales, three mangroves *Hibiscus tiliaceus*, *Heritiera littoralis* and *Thespesia populnea*, together with Huerteales, Brassicales species were clustered as neighbor orders. The relationship of genus within the families of Malvales is in accord with chloroplast reports of *Aquilaria sinensis* [19] and *Heritiera angustata* [20], and additionally confirmed that the semi-mangrove *Thespesia populnea* is close to *Gossypium* plants. For Malpighiales, an order of astonishing morphological and ecological diversity, the phylogenetic relationship of different families, especially Linaceae, is less resolved. With the exception of grouping of families Linaceae and Euphorbiaceae, our phylogenetic tree is largely consistent with a 2012 study, which employed 82 plastid genes from 58 species, representing 39 of the 42 families of Malpighiales [21]. We found that Euphorbiaceae constituted a single branch, while Rhizophoraceae (including four mangroves: *Bruguiera sexangular*, *Kandelia obovata*, *Rhizophora stylosa*, *Ceriops tagal*) is neighboring Erythroxylaceae and Clusiaceae, and Linaceae is a sister to Chrysobalanaceae and Malpighiaceae. This relationship is supported by a very recent study [22]. Finally, our phylogenetic tree supported the sister relationship between the mangrove *Pongamia pinnata* included order Fabales and Rosales, Cucurbitales and Fagales. Species that from one order are clustered together, and their chloroplast genome relations among different orders are almost consistent with the public resource [23]. Our study used the whole chloroplast genomic information to infer the evolutionary relationships, and these phylogenetic trees using multiple methods not only provide genome-scale support for the 14 mangroves but also offer insights for the positional relationship of angiosperms in several orders and families.

### Genomic synteny and divergence

Gene orientation was assessed by synteny analysis among the 71 species based on their phylogenetic relationships, revealing conserved gene structure and distribution in plant chloroplast genomes, and only one mangrove *Heritiera littoralis* suffered gene re-arrangement in order Malvales (Figure 2). According to the synteny distribution, a notable rearrangement, from 8,109 bp to 3,3498 bp, occurred in *Heritiera littoralis* comparing to *Hibiscus tiliaceus* and *Thespesia populnea*. This region encodes 16 genes: *trnC-GCA*, *petN*, *psbM*, *trnD-GUC*, *trnY-GUA*, *trnE-UUC*, *rpoB*, *rpoC1*, *rpoC2*, *rps2*, *atpI*, *atpH*, *atpF*, *atpA*, *trnR-UCU*, and *trnS-CGA*. To measure the genetic divergence of the mangrove chloroplast genomes, the sequence similarity was assessed in the orders. We found that similarity of coding regions is much higher than the intergenic regions, and the tRNA and rRNA genes are almost identical in all the species (Figure S6). Another phenomenon observed in all six orders, is that SSC regions and LSC have higher divergence than IR regions, indicating the IR is more conserved than single copy sequences (Figure 3). This pattern is also been reported in other plants [24–26].

**Figure 2.**
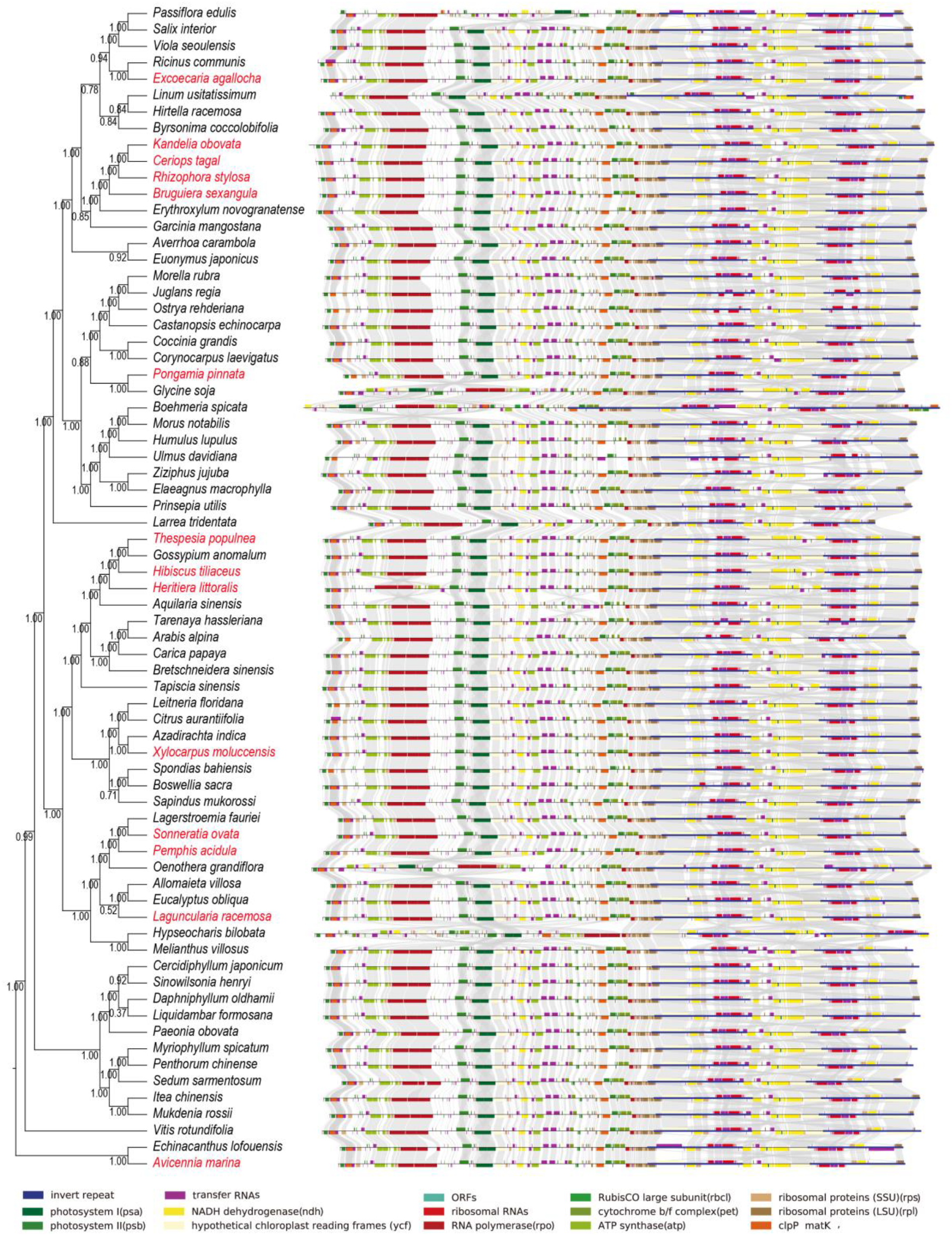
The gene orders’ comparison of whole chloroplast genome between mangroves (red) and the other species in phylogenetic tree.

**Figure 3.**
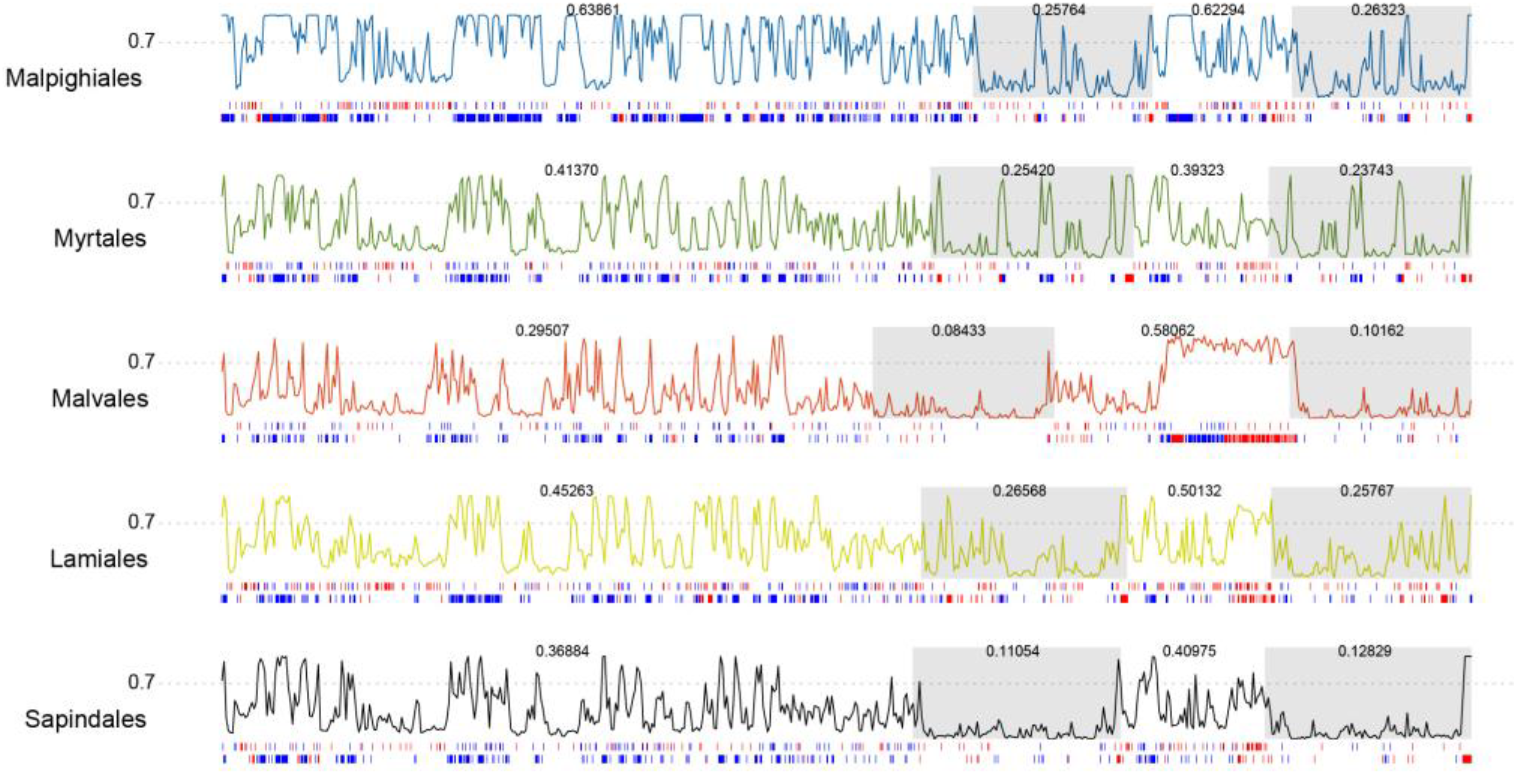
The distribution of divergence (polylines) within orders of Malpighiales, Myrtales, Malvales, Lamiales, and Sapindales. The IR regions were marked with grey background. The average variation values of each region were showed above the lines. Two bars at the bottom of the line shows SNP density and InDel in gene regions (red) and intergenic regions(blue).

### Analysis of selective pressures

The nonsynonymous substitutions and synonymous substitution ratio (Ka/Ks) were calculated for 73 chloroplast genes in 14 mangroves and their close species (see Methods). Branches with Ka/Ks values above 1.0 are under positive selection and are candidates for functional adaptations, while values less than 1.0 is indicative of negative (purifying) selection [27]. As shown in Figure 4, the Ka/Ks values of most pairs (within species and between orders) was consistently less than 1.0, suggestive of evolutionary pressure to maintain gene function. For instance, genes associated with Photosystems (*psaA*, *psaB*, *psaC*, *psaJ*, *psbA*, *psbC*, *psbD*, *psbE*, *psbF*, *psbH*, *psbL*, *psbM*, and *psbT*), the Cytochrome b/f complex (*petA*, *petB*, *petD*, *petG*, and *petN*), and some ATP synthase (*atpA*, *atpB*, *atpH*, and *atpI*) in all species had a Ka/Ks value close to 0. Genes encoding some ATP synthase (*atpA*, *atpE*, and *atpF*, and *atpI*), NADH dehydrogenase (*ndhA*, *ndhB*, *ndhC*, *ndhD*, *ndhE*, *ndhF*, *ndhG*, *ndhH*, *ndhI*, *ndhJ*, and *ndhK*), Ribosomal proteins (*rps2*, *rps3*, *rps4*, *rps8*, *rps11*, *rps12*, *rps14*, *rps15*, *rps16*, *rps18*, and *rps19*), and RNA polymerase (*rpoA*, *rpoB*, *rpoC1*, and *rpoC2*), also had low Ka/Ks values (mostly between 0~0.5). Six genes (*petL*, *psaI*, *rpl23*, *rpl36*, *rps7*, and *ycf1*) have Ka/Ks value greater than 1.0 in at least one mangrove species and confirmed by two methods (Figure 4, Table S4). These genes could be classified into four groups according to their function: subunits of cytochrome (*petL*), subunits of photosystems (*psaI*), subunits of ribosome (*rps7*, *rpl23*, and *rpl36*), and unclassified gene (*ycf1*). The gene *petL* is a component of the cytochrome b6/f complex required for photosynthesis. In this study, the Ka/Ks value of *petL* in *Sonneratia ovata* and *Pemphis acidula* from Myrtales was ~1.5, and we found the values in this order ranges slightly larger than in Malvales and Sapindales (Figure 4). While a member of Phostosystem I (PSI), *psaI*, was observed that has a Ka/Ks value of 1.22 in *Laguncularia racemose*, suggesting potential positive selection in this species. According to the distribution, this gene has a wide range of Ka/Ks values from 0.2 to 1.5, especially in Malpighiales, Myrtales, and Sapindales, which may reflect the various adaptation in different clades. Although research of *psaI* in tobacco found a role for *psaI* in stabilizing PSI during leaf senescence [28], the function of the protein remains unknown in most plant species. Among the LSU and SSU genes, *rpl23* and *rpl36* (LSU) and rps7 (SSU) had Ka/Ks values greater than 1.0 in *Ceriops tagal*, *Excoecaria agallocha*, *Heritiera littoralis*, *Hibiscus tiliaceus*, *Kandelia obovata*, *Pemphis acidula*, *Sonneratia ovata*, *Thespesia populnea*, and *Xylocarpus moluccensis*. In a previous study [29], *rpl23* was reported to be essential for plants and loss of *rpl36* result in severe morphological aberrations, low translational efficiency, poor photoautotrophic growth [29]. As an rRNA binding protein, *rps7* plays a role in the regulation of chloroplast translation [30]. However, the experimental research on the specific function of *rps7* is limited at present. We speculate that relaxed neutral selection (Ka/Ks value around 1.0) and potential positive selection (Ka/Ks value greater than 1.0) of *rpl23*, *rpl36*, and *rps7* in various species here reflects the highly diverse environments and adaptions of plants (Table S4). For gene *ycf1*, it is one of the largest genes in chloroplast genome and usually has two functional copies [22,25], or one of the two copies was evolved to be a pseudogene in some plants [19,31–33] as well as in 11 mangroves (*Avicennia marina*, *Xylocarpus moluccensis*, *Hibiscus tiliaceus*, *Excoecaria agallocha*, *Bruguiera sexangula*, *Kandelia obovata*, *Rhizophora stylosa*, *Ceriops tagal*, *Sonneratia ovata*, *Pemphis acidula*, *Laguncularia racemose*). In this study, most species in Malpighiales including one mangrove *Excoecaria agallocha*, have *ycf1* Ka/Ks values around or greater than 1.0, while about 0.4 in Malvales, Myrtales and Sapindales, and about 0.6 in Fabales and Lamiales, showing the diverse evolve rates in different species (Table S4, Figure 4). This is consistent with the lower similarity of the sequences in gene *ycf1* (Figure S6). In summary, the Ka/Ks values here provide comparative results for detection of selective pressure in mangrove chloroplasts. The genes that involve photosynthesis and energy metabolism for chloroplasts are conserved with very low Ka/Ks ratio, and this kind of evolutionary pattern has been generally observed in other plant chloroplasts [22,32]. The potential positive selection or relax pressure of *ycf1*, *rpl23*, *rps7*, *rpl36*, *petL*, *psaI* were also detected in mangroves, indicating their fast evolve rate and divergence in different species in spite of its ambiguous function in chloroplast genomes.

**Figure 4.**
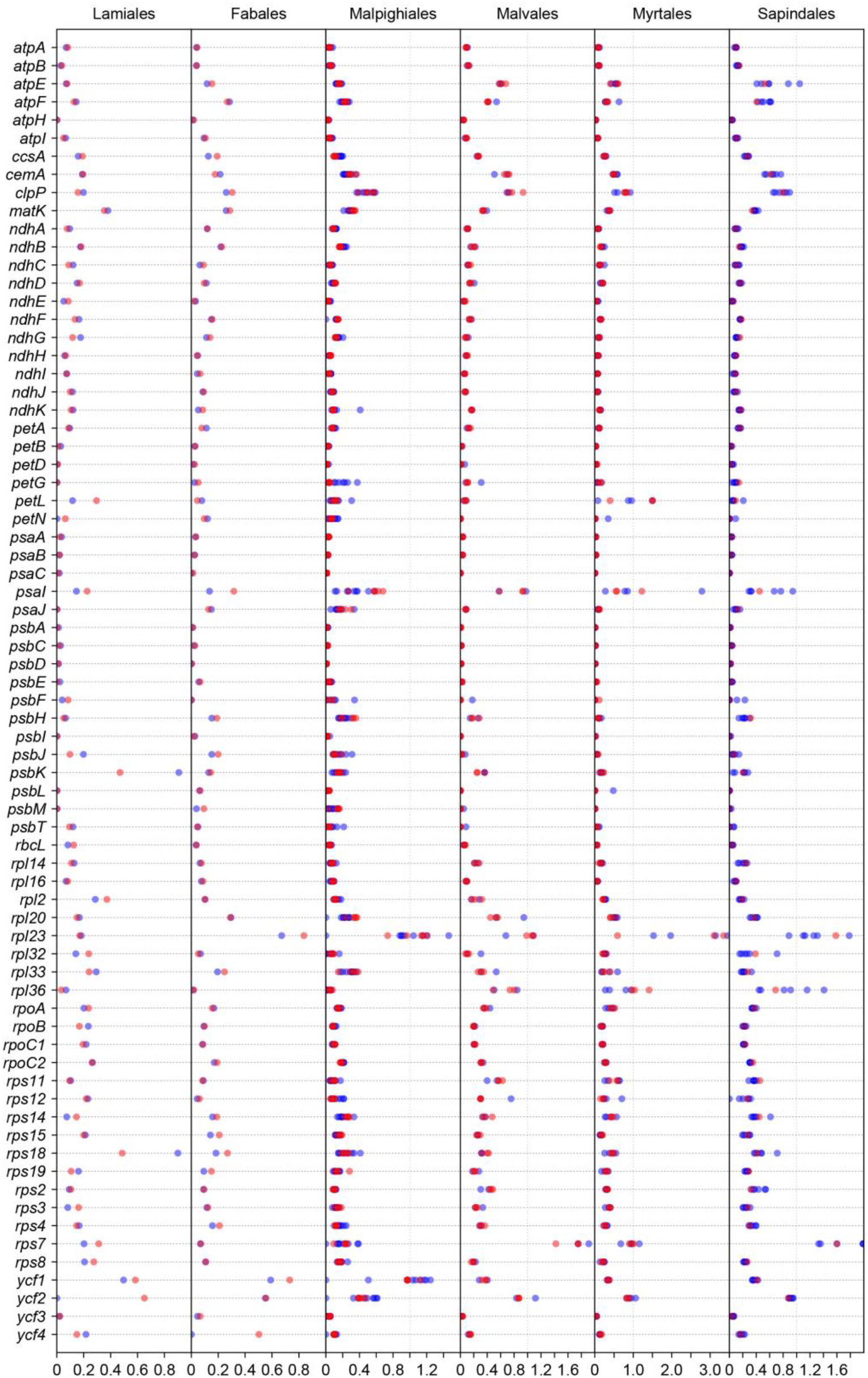
The Ka/Ks values of 73 protein-coding genes of the mangroves species and their sister species in six orders (Lamiales, Fabales, Malpighiales, Malvales, Myrtales, Sapindales). Circles in red represent mangrove, while blue ones represent non-mangrove species.

## Methods

### Chloroplast genome sequencing, assembly and annotation

Fresh leaves from mangroves were collected in Guangzhou, China and DNA were extracted and stored in BGI-Qingdao and sequenced on BGISEQ500 platform. Considering the small genome size of a chloroplast, about five hundred million pair-end sequencing data was randomly used from each species for further assembly. These data were processed by MITObim v1.9 [34] as initial assembly using closest reference-referred assembly strategy. Total length of each chloroplast were estimated by SPAdes v3.13.0 [35]. Then NOVOPlasty v2.7.2 [36] was performed to assemble a complete genome based on the initial assembly information and genome size information. Finally, manual orientation and connection was further carried out to acquire circular sequences. The data that support the findings of this study have been deposited in the CNSA (https://db.cngb.org/cnsa/) of CNGBdb with accession number CNP0000567.

Chloroplast genes including protein coding, rRNA and tRNA were predicted and annotated on the website tool GeSeq [37] using MPI-MP chloroplast references option. The identity cutoff for protein and rRNA searching were set with 60 and 85 respectively. The ARAGORN v1.2.38 was chosen for tRNA annotation. The information of genes along the whole chloroplast genome were visualized by OGDRAW [38] with default settings. The IR (Inverted Repeat) boundaries in chloroplast sequences were identified by aligning the whole sequence with itself using BLAST (-p blastn -m 8 -F F -e 1). The results that with completely reverse alignment between two regions of equal length were selected and manually checked as final Inverted Repeat regions. The simple sequence repeat (SSR) that with 1-6bp were detected using MISA program [39]. The minimum repeats were set with 10 for mononucleotides, 5 for dinucleotide, 4 for trinucleotide, 3 for tetranucleotide, 3 for pentanucleotide and 3 for hexanucleotide.

### Phylogenetic analysis

A total of 41 conserved genes (*atpA*, *atpB*, *atpE*, *atpH*, *atpI*, *ccsA*, *cemA*, *matK*, *ndhA*, *ndhC*, *ndhG*, *ndhI*, *ndhJ*, *petA*, *petN*, *psaA*, *psaB*, *psaC*, *psaJ*, *psbA*, *psbC*, *psbD*, *psbE*, *psbF*, *psbH*, *psbI*, *psbJ*, *psbT*, *rpl14*, *rpl2*, *rpoA*, *rpoB*, *rps11*, *rps14*, *rps15*, *rps19*, *rps2*, *rps3*, *rps4*, *rps8*, *ycf3*) which exists in all the 71 species’ chloroplast genomes were used to construct a reliable phylogenetic tree (species were list in Table S2). The coding sequences were aligned by MAFFT (v7.407) [40] with “--auto --adjustdirection” setting. Based on the global alignments, ML phylogenetic tree was constructed using RAxML program (version 8.2.12) [41] with GTRGAMMAI model, and iqtree [42] with best-fit partition model [43]. The bootstrap for ML trees was set with number 1000. The BI trees were constructed using MrBayes (version 3.2.7) [44] with GTR+GAMMA model. Trees were assessed by CONSEL (version 1.20) [45] and iqtree [42].

### Ka/Ks calculation

The non-synonymous (Ka) and synonymous (Ks) substitution ratio (Ka/Ks) of genes in 14 mangroves, as well as species from Lamiales (*Echinacanthus lofouensis*: NC_035876.1) Fabales (*Glycine soja*: NC_022868.1), Malpighiales (*Hirtella racemosa*: NC_024060.1, *Salix interior*: NC_024681.1, *Viola seoulensis*: NC_026986.1, *Ricinus communis*: NC_016736.1, *Erythroxylum novogranatense*: NC_030601.1, *Passiflora edulis*: NC_034285.1, *Garcinia mangostana*: NC_036341.1, *Linum usitatissimum*: NC_036356.1, *Byrsonima coccolobifolia*: NC_037191.1), Malvales (Gossypium anomalum: NC_023213.1, *Aquilaria sinensis*: NC_029243.1), Myrtales (*Eucalyptus obliqua*: NC_022378.1, *Oenothera grandiflora*: NC_029211.1, *Lagerstroemia fauriei*: NC_029808.1, *Allomaieta villosa*: NC_031875.1), and Sapindales (*Azadirachta indica*: NC_023792.1, *Sapindus mukorossi*: NC_025554.1, *Citrus aurantiifolia*: NC_024929.1, *Boswellia sacra*: NC_029420.1, *Leitneria floridana*: NC_030482.1, *Spondias bahiensis*: NC_030526.1) were calculated. As these orders cover asteridsI (Lamiales), rosids I (Fabales, Malpighiales) and rosids II (Malvales, Myrtales, and Sapindales) of core eudicots, we chose three species from order Asterales (NC_023109.1) in asterids I clade, order Zygophyllales (NC_028023.1) in rosid I and order Geraniales (NC_023260.1) in rosid II as an outgroup for calculation. Genes from each chloroplast genome were compared with the corresponding outgroup using MAFFT (v7.407) [40] to generate pairwise alignment sequences. Then Ka/Ks values were calculated by KaKs Calculator program [46] and the target genes were re-calculated by PAML [47].

### Divergence calculation

Whole chloroplast sequences from same order including mangroves were used to calculate divergence. The whole genome sequences of Lamiales species (*Sesamum indicum*: NC_016433.2, *Lindenbergia philippensis*: NC_022859.1, *Ajuga reptans*: NC_023102.1, *Hesperelaea palmeri*: NC_025787.1, *Scrophularia takesimensis*: NC_026202.1, *Tanaecium tetragonolobum*: NC_027955.1, *Erythranthe lutea*: NC_030212.1, *Paulownia coreana*: NC_031435.1, *Haberlea rhodopensis*: NC_031852.1, *Aloysia citrodora*: NC_034695.1, *Echinacanthus lofouensis*: NC_035876.1, and one mangrove *Avicennia marina*), Malpighiales (*Byrsonima coccolobifolia*: NC_037191.1, *Erythroxylum novogranatense*: NC_030601.1, *Garcinia mangostana*: NC_036341.1, *Hirtella racemosa*: NC_024060.1, *Ricinus communis*: NC_016736.1, *Salix interior*: NC_024681.1, *Viola seoulensis*: NC_026986.1, and mangroves *Bruguiera sexangula*, Ceriops tagal, *Excoecaria agallocha*, *Kandelia obovata*, *Rhizophora stylosa*), Myrtales (*Allomaieta villosa*: NC_031875.1, *Eucalyptus obliqua*: NC_022378.1, *Lagerstroemia fauriei*: NC_029808.1, and mangroves *Laguncularia racemosa*, *Pemphis acidula*, *Sonneratia ovata*), Sapindales (*Azadirachta indica*: NC_023792.1, *Boswellia sacra*: NC_029420.1, *Citrus aurantiifolia*: NC_024929.1, *Leitneria floridana*: NC_030482.1, *Sapindus mukorossi*: NC_025554.1, *Spondias bahiensis*: NC_030526.1, and mangrove *Xylocarpus moluccensis*), Malvales (*Gossypium arboretum*: NC_016712.1, *Daphne kiusiana*: NC_035896.1, and mangroves *Hibiscus tiliaceus*, *Thespesia populnea*) were aligned with MAFFT (v7.407) [40] program within orders respectively then SNP and InDel sites were calculated in 200bp windows. Genomic comparison and similarities were performed by mVISTA [48].

### Conclusions

This study is the first large-scale report of whole-genome sequence of chloroplasts of mangroves from six orders (five rosids and one asterid). They have identical genomic IR regions with similar length, GC content, and gene number. The mangroves *Avicennia marina*, the only asterids plant in this study, and *Heritiera littoralis*, one rosids plants, have the translation initiation factor gene *infA*. The genomic structure of mangrove chloroplasts was highly similar. In the 14 mangroves, only *Heritiera littoralis* contained an inversion event in the LSC region. The phylogenetic tree constructed based on conserved genes in 71 species encompasses most dicot families and represents the first complete phylogenetic analysis of mangrove plants where rosids are included. Non-synonymous and synonymous substitutions (Ka/Ks) analysis revealed evidence of positive selection of six genes in ten mangroves and showed varied evolutionary rates among genes in different chloroplast genomes. In conclusion, we assembled new 14 complete chloroplast genomes from diverse mangroves and analyzed their genomic characteristics, providing a useful resource for future studies aiming to resolve the ecology and evolution of these enigmatic group of plants.

